# Improving replicability in single-cell RNA-Seq cell type discovery with Dune

**DOI:** 10.1101/2020.03.03.974220

**Authors:** Hector Roux de Bézieux, Kelly Street, Stephan Fischer, Koen Van den Berge, Rebecca Chance, Davide Risso, Jesse Gillis, John Ngai, Elizabeth Purdom, Sandrine Dudoit

## Abstract

Single-cell transcriptome sequencing (scRNA-Seq) has allowed many new types of investigations at unprecedented and unique levels of resolution. Among the primary goals of scRNA-Seq is the classification of cells into potentially novel cell types. Many approaches build on the existing clustering literature to develop tools specific to single-cell applications. However, almost all of these methods rely on heuristics or user-supplied parameters to control the number of clusters identified. This affects both the resolution of the clusters within the original dataset as well as their replicability across datasets. While many recommendations exist to select these tuning parameters, most of them are quite ad hoc. In general, there is little assurance that any given set of parameters will represent an optimal choice in the ever-present trade-off between cluster resolution and replicability. For instance, it may be the case that another set of parameters will result in more clusters that are also more replicable, or in fewer clusters that are also less replicable.

Here, we propose a new method called `Dune` for optimizing the trade-off between the resolution of the clusters and their replicability across datasets. Our method takes as input a set of clustering results on a single dataset, derived from any set of clustering algorithms and associated tuning parameters, and iteratively merges clusters within partitions in order to maximize their concordance between partitions. As demonstrated on a variety of scRNA-Seq datasets from different platforms, `Dune` outperforms existing techniques, that rely on hierarchical merging for reducing the number of clusters, in terms of replicability of the resultant merged clusters. It provides an objective approach for identifying replicable consensus clusters most likely to represent common biological features across multiple datasets.

---

Improvements in single-cell transcriptome sequencing (scRNA-Seq) over the last decade have allowed the characterization of gene expression in collections of thousands to hundreds of thousands of cells. While datasets have grown in size by several orders of magnitude, cell type identification remains a primary step in the analysis process [1]. We will focus here on unsupervised clustering, which can be broadly defined as partitioning observations into clusters based on a set of features, without using any prior knowledge on the groupings. In the scRNA-Seq context, clustering aims to identify groups of cells that are defined by a unique and consistent transcriptomic signature. Such groups of cells can represent both transient features, such as cellular states, or more permanent features, such as celullar types.

Many clustering algorithms have been proposed for scRNA-Seq, most of these being adaptations from the clustering literature at large. Popular methods include `SC3` [2], `Seurat` [3], and `Monocle` [4]. However, clustering remains a complex task. Kiselev et al. [5] outlined the various challenges – both biological and computational – of this step, including technical noise, biological heterogeneity, and the impact of tuning parameters for the clustering algorithms. In particular, obtaining replicable clusters can be difficult. In this work, we declare clusters as replicable if running the exact same clustering algorithm on a related dataset yields similar clusters. Duò et al. [6] offers a recent review and benchmark of some scRNA-Seq clustering algorithms, identifying `SC3` and `Seurat` as the best methods overall. The selection of tuning parameters, however, remains an open question. While some methods, `SC3` for example, provide a way to estimate the optimal value of its main tuning parameter, most do not, leaving the choice to the user. Consensus methods try to bypass this issue [2, 7], but they also rely on meta-parameters which can still have substantial impact on the results.

The aforementioned clustering algorithms identify a pre-specified number of clusters either directly, as in *k*-means, or indirectly, through another tuning parameter. They rely on the assumption that there is only one relevant level of clustering resolution, i.e., an optimal number of clusters, in the dataset. We argue that this is often not the case, since cell types usually have a hierarchy. For example, Tasic et al. [8] propose a tree structure for the mouse anterolateral motor (ALM) and primary visual (VISp) cortical areas. At the higher levels, cells can be clustered as neurons and non-neurons. Then, neurons can be further split into GABAergic and glutamatergic neurons and so on and so forth. This hierarchical structure means that the concept of an “optimal” number of clusters is not appropriate. Instead, many datasets can be better characterized with ever-finer levels of resolution. At the higher levels, cells are grouped into large clusters that are quite coarse, but are easily identifiable and very replicable across datasets. As the resolution increases, there are more and more clusters, but these are less and less certain, meaning that they are less likely to represent real biological cell types and more likely to be reflecting over-partitioning (cf. overfitting) of the data or the presence of transient states. This resolution-replicabilty trade-off is not obvious to quantify and is heavily dataset-dependent: it is not only influenced by the biological setting under study and its complexity, but also depends on technical properties of the data, such as sequencing depth and number of cells [1].

By far the most common method to establish a hierarchy for pre-defined clusters is agglomerative hierarchical clustering, a bottom-up method in which clusters are merged one-by-one until they are all merged into a single cluster. This procedure yields a tree structure linking clusters that are merged together. The tree can also be defined by merging clusters according to the fraction of differentially expressed (DE) genes between them [7, 8]. While several extensive benchmarks of clustering methods have been proposed [6, 9], these only focus on the resulting partitions rather than the full hierarchical structure. Zappia and Oshlack [10] proposed a representation of clustering trees to visually describe hierarchies but this type of analysis heavily relies on user-supervision.

Here, we present `Dune`, a method that aims to reconciles multiple clustering results and extract the common structure that they all identify. `Dune` takes as input a set of clustering results (i.e., results from a variety of clustering algorithms and associated tuning parameters applied to a given dataset) and produces hierarchies of clusters by merging clusters within each partition using information borrowed from the other partitions. While different clustering algorithms run with different tuning parameters will naturally provide discrepant clusters, all good clustering methods should be able to identify a common higher-level clustering that is robust to the choice of tuning parameters. `Dune` identifies this common higher level of resolution shared by all methods without requiring any tuning by the user. Examining this level can provide both useful biological insight and help to compare various clustering methods.

In this manuscript, we first introduce the `Dune` algorithm. Then, using a variety of scRNA-Seq and snRNA-Seq datasets from different sequencing platforms, we show that `Dune` outperforms agglomerative merging methods in navigating the trade-off between resolution and replicability and in identifying gold-standard high-level clusterings. Finally, we assess `Dune`’s robustness to poor inputs and to sample size.

## Results

### The Dune algorithm

The `Dune` algorithm is a general framework that increases the agreement between different clusterings of the same dataset through iterative merging. It takes as input *R* sets of clustering results, generally produced from running *R* clustering algorithms (or the same algorithm with different tuning parameter values) on the same dataset. An example can be seen in Figure 1a, where a small subset of the AIBS snRNA-Smart dataset [11] (see the “Methods, Case Studies” section) is used to demonstrate some of the main concepts underlying `Dune`. The first row displays three examples of clusterings (i.e., sets of cluster labels) produced by three different clustering algorithms applied to the same dataset, reduced to two dimensions using t-SNE[12–14]. All three methods identify similar, but not identical clusters. Indeed, the algorithms output partitions with different levels of resolution. For example, `Monocle` splits the bottom region (in reduced dimension) into two clusters, while the other two methods find three clusters. Likewise, `Monocle` and `SC3` find two clusters in the top region, while `Seurat` only finds one. These differences can be displayed using confusion matrices (second row of Figure 1a), where the overlap between two clusters from any pair of clusterings is displayed both in terms of the number of cells in the intersection and by the Jaccard index (i.e., the cardinality of the intersection of the two clusters over the cardinality of their union; [15]). Rows and columns are ordered so as to maximize, as much as possible, the sum of the diagonal entries. Confusion matrices can be further summarized using the adjusted Rand index (ARI). The ARI [16, 17] is a commonly used measure for the agreement between two sets of clustering labels, see the “Methods, ARI” section for more details. As can be seen in the confusion matrices, `SC3` and `Seurat` have the highest level of agreement. Indeed, this is also reflected in the fact that they have the highest ARI of any pair.

**Figure 1:**
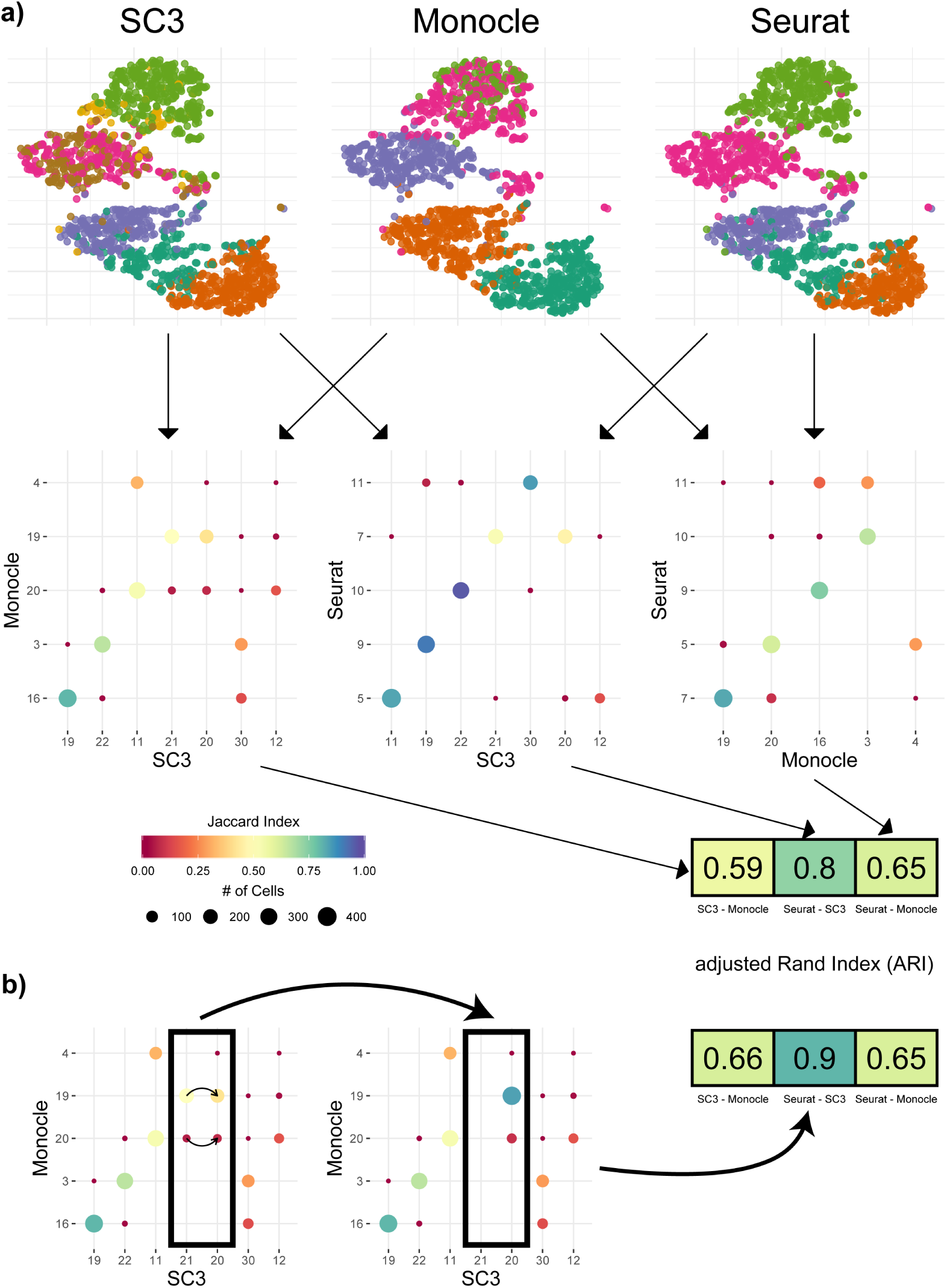
Measuring and improving the concordance between clusterings. We used a subset of the AIBS snRNA-Smart dataset as an example. Panel **a.** `SC3`, `Monocle`, and `Seurat` were run on the dataset and their results are displayed using scatterplots of the first two t-SNE components, where the color of the plotting symbol corresponds to the cluster label. Each pair of clusterings was then compared using a confusion matrix, resulting in three such matrices. For a pair of clusterings/partitions, a confusion matrix is a contingency table, where each entry corresponds to the number of observations in both a cluster from the first partition and a cluster from the second. The size of the dot represents the number of observations in both clusters and the color corresponds to the Jaccard index. Each confusion matrix produces one ARI value. Panel **b.** Merging clusters 20 and 21 from `SC3` into one cluster changes the confusion matrix and increases the ARI.

`Dune` merges together the clusters within each of the *R* partitions so that the *R* clustering results more closely match each other. An example of the merging is displayed in Figure 1. Clusters 20 and 21 from `SC3` are merged together, resulting in one larger cluster named 20. Doing so increases the agreement between `SC3` and `Monocle` in the confusion matrix, as reflected by an increase in ARI from 0.59 to 0.66. This merge also improves the ARI between `SC3` and `Seurat` (from 0.8 to 0.9) and hence increases the overall agreement between the three clusterings. This is the main idea behind `Dune`. Specifically, `Dune` performs an iterative search, where, at each iteration, it identifies the partition and pair of clusters within this partition that, when merged, most improve the average of the adjusted Rand index over all pairs of clusterings (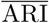). Thus, the `Dune` algorithm can be viewed as an iterative algorithm for maximizing the average pairwise ARI of a collection of clustering results. A more formal definition of the algorithm is provided in the “Methods, Dune” section.

We demonstrate how the `Dune` algorithm works in Figure 2, using the AIBS scRNA-Smart dataset, a scRNA-Seq dataset of 6,300 mouse brain cells further described in the “Methods, Case Studies” section. For this example, we ran `SC3`, `Seurat`, and `Monocle` to obtain our initial clustering results for input into `Dune` (*R* = 3). Figure 2a displays the confusion matrix for a pair of clusterings (`SC3` and `Monocle`) before any merging and Figure 2b displays a pseudocolor image of the matrix of all pairwise ARIs for the three clusterings before any merging. The overlap between the three methods is moderate. Indeed, the pairwise ARIs vary between 0.55 and 0.68 in Fig. 2b. However, as can be seen in the confusion matrix, the clusterings do capture a shared underlying structure, which will serve as grounding for the `Dune` merging. Figure 2d shows the confusion matrix for the same two partitions as in 2a, after merging with `Dune`. We can see that we have, by design, fewer clusters in both partitions, but also that the concordance between the two partitions is greatly improved (as indicated by the color of the plotting symbols, which represents the Jaccard Index). This is further evidenced in Figure 2e, where the pairwise ARIs between the three partitions are displayed. The average ARI after all merging steps increased from ∼ 0.6 to ∼ 0.89. Figures 2c and 2f demonstrate the evolution of the average ARI and of the number of clusters per partition through the `Dune` merging process. At each step, we merge the pair of clusters that leads to the greatest increase in average ARI. Hence, at each step, the average ARI increases (Fig. 2c) and the number of clusters in one of the partitions decreases by one (2f). The final partitions are achieved when the average ARI can no longer be improved.

**Figure 2:**
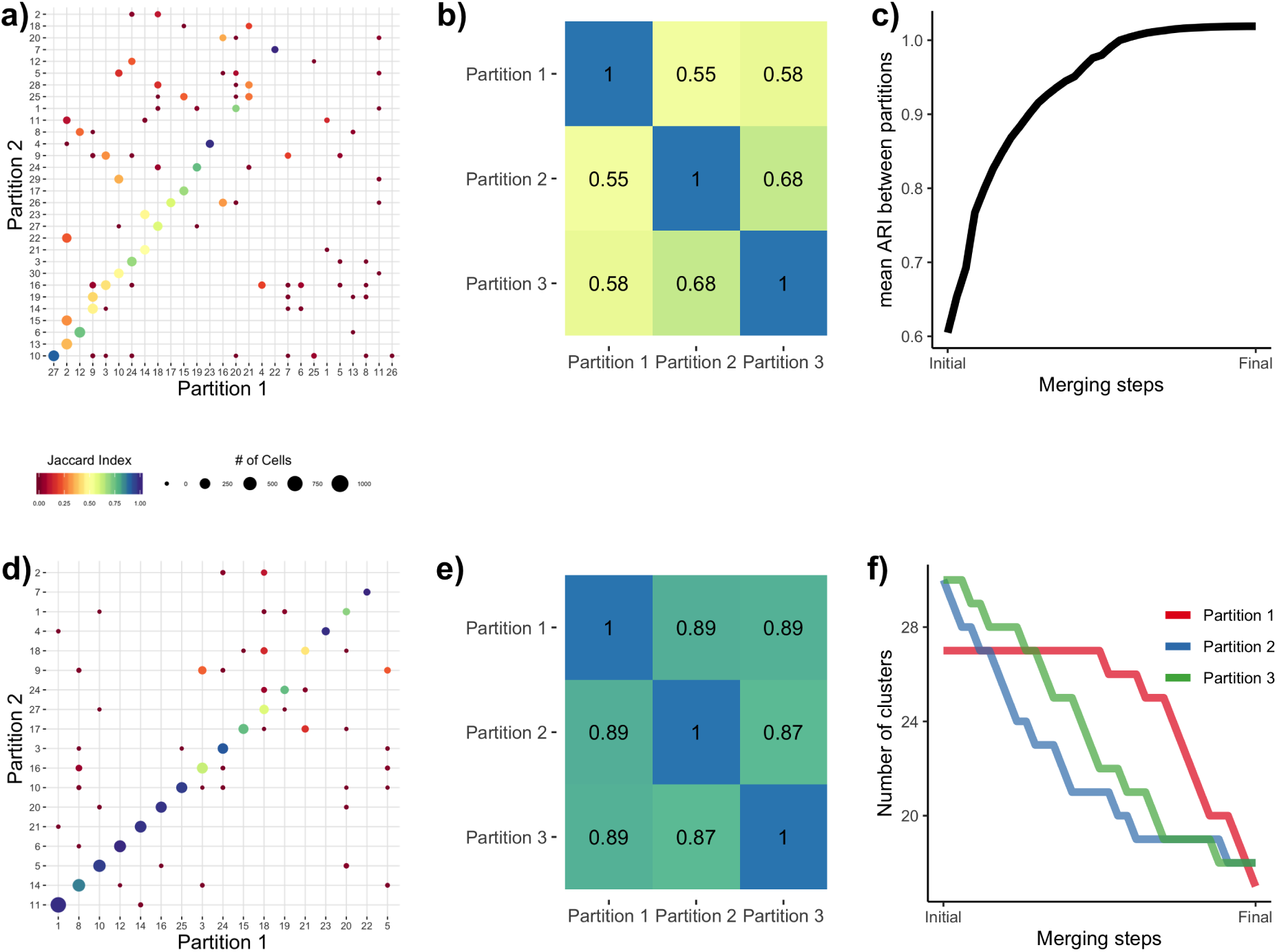
Illustrating Dune on a dataset with three sets of clusters. We used the AIBS scRNA-Smart dataset [11] as an example. Before any merging, the sets of cluster labels – or partitions – resulting from running `SC3`, `Seurat`, and `Monocle` have a moderate agreement. Panel **a** displays the confusion matrix between two of the partitions, where each entry corresponds to the number of observations in both a cluster from Partition 1 and a cluster from Partition 2. The confusion matrix shows that while many cells are clustered in similar clusters, i.e., along the main diagonal, many others are not. This can be summarized by the ARI between Partitions 1 and 2. Panel **b** displays a pseudocolor image of the matrix of all pairwise ARIs between the three partitions. Panel **c** illustrates that the average ARI between partitions increases as pairs of clusters are merged when applying `Dune`. After running `Dune`, the confusion matrix in Panel **d** and the pairwise ARI matrix in Panel **e** both show that the partitions are indeed more similar. Panel **f** shows that, at each merging step, the number of clusters in one of the partitions is decreased by one, in `Dune`’s greedy procedure to improve the average ARI by merging pairs of clusters.

In the following sections, we evaluate `Dune` and compare it to two hierarchical tree merging methods, using four datasets: two mouse brain datasets from the Allen Institute *** HRB: waiting for main paper and two human pancreas datasets [18, 19]. We then discuss the value of `Dune`’s stopping rule. Finally, we investigate the stability of the `Dune` algorithm to the clustering inputs and the sample size.

### Dune outperforms other methods in recovering known biological subtypes

To evaluate `Dune`, we first considered how well the resulting merged clusters compare to known biological subtypes. We used the output of `Dune` on the *R* = 3 clustering methods (namely, `SC3`, `Seurat`, and `Monocle`) applied to the AIBS scRNA-Smart dataset, as described above. For this dataset, we treated the labels from the original publication as the gold standard. At each merge (i.e., iteration), we computed the ARI between the the known subtypes and the `Dune` clusters. Figure 3a displays the ARI evolution for the clusters from `SC3` as they are merged with `Dune` (blue curve). As merging occurs, the resolution (i.e., number of clusters) decreases and the ARI with the known cell subtypes increases. The entire ARI curve can be summarized by computing the the area under it, referred to herein as the area under the ARI curve (AUARIC), as depicted in Figure 3b.

**Figure 3:**
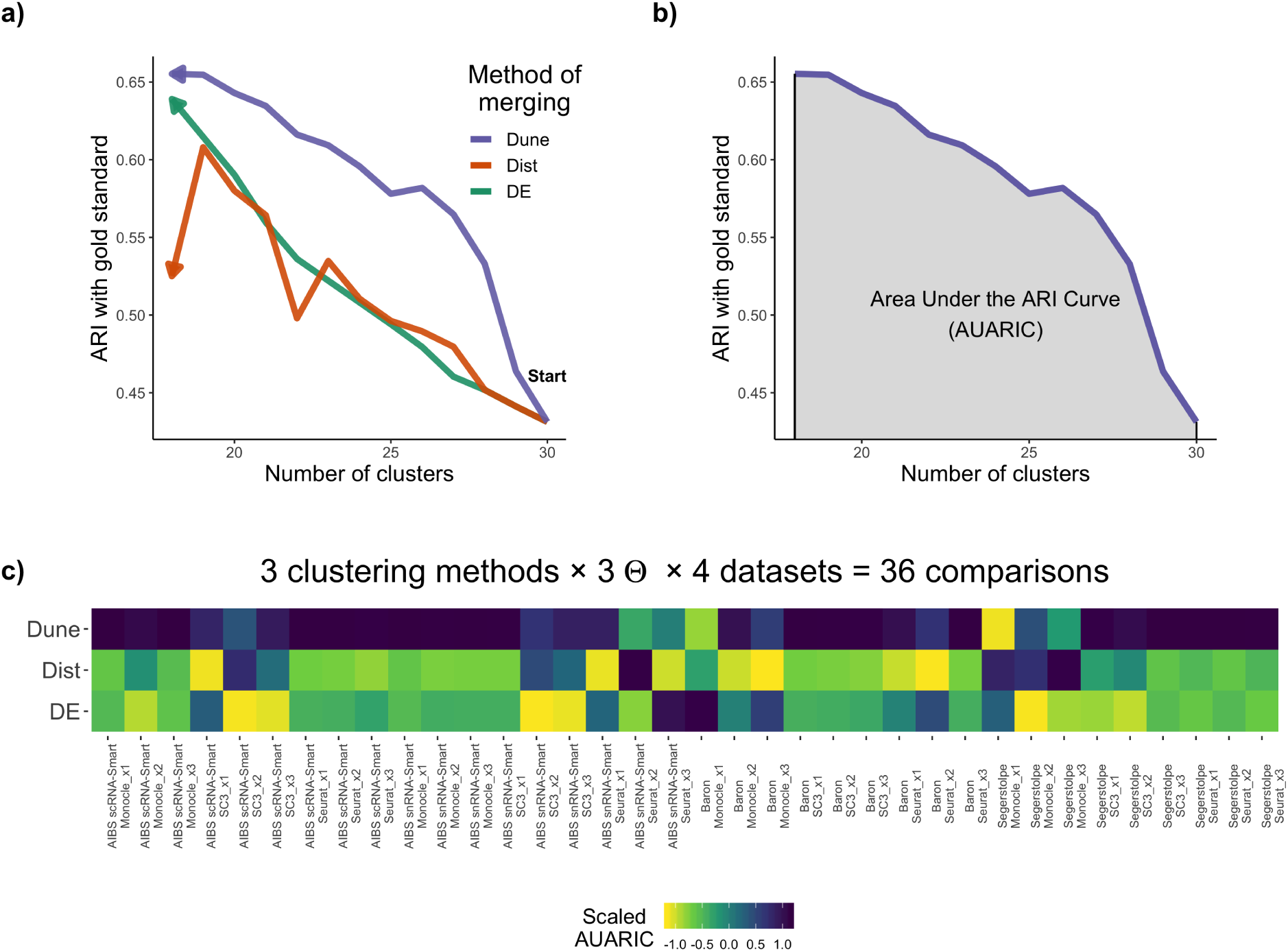
Dune outperforms other methods in recovering known biological subtypes. Panel **a.** `SC3` was run on the AIBS scRNA-Smart dataset for *θ*_*sc*3_ = 0 and merged down with either `DE`, `Dist`, or `Dune` (with *θ*_*Monocle*_ = 45 and *θ*_*Seurat*_ = 1.2, for `Dune`). The ARI with the labels from the original publication, treated as gold standard, was computed at each step of all three merging procedures. Panel **b.** For each merging method from **a**, the area under the ARI curve (AUARIC) was computed. This was repeated for three clustering methods, each with three different values of their respective tuning parameter *θ*, and four datasets. The resulting 36 AUARIC are displayed in the pseudocolor image of Panel **c**. The AUARIC values are scaled to have a column mean of zero and column variance of 1. This was done to make AUARIC values comparable across datasets, clustering methods, and parameter values, since the AUARIC can have different scales across scenarios.

We compared the performance of `Dune` to other methods of merging, referred to as `Dist` and `DE` (red and green curves in Figure 3a, respectively). Both are hierarchical methods, that start by building a tree between the clusters. The `Dist` method then merges clusters in a bottow-up manner, starting with the two clusters that are closest in the tree and then iteratively until all clusters are merged. The second approach, `DE`, follows the method implemented in `RSEC` and merges clusters bottom-up based on the percentage of DE genes between clusters. It uses the `limma` package [20], where a gene is declared DE if its nominal false discovery rate (FDR) adjusted *p*-value is below 0.05 [21]. Pairs of clusters with less than a certain fraction of DE genes are merged. Increasing this threshold from 0 to 1 leads to an iterative merging procedure. More details on these two procedures can be found in the Method section.

In Figure 3a, we see that `Dune` consistently outperformed the other two integration methods in terms of concordance with BICCN-curated clusters throughout the merging process and therefore also in term of AUARIC. We note that while `Dune` stops merging when the average ARI can no longer be improved, the hierarchical merging procedures have no meaningful stopping point and continue merging until only one cluster is left. To provide a reasonable stopping point, we stopped the other methods when merging no longer improves the ARI, similar to the requirement of `Dune`, which means we did not penalize the other methods for not providing a natural stopping point. For each merging method, we computed an area under its ARI curve (AUARIC), as depicted in Figure 3b for the merging of the `SC3` clusters of the AIBS scRNA-Smart dataset using `Dune`.

Figure 3c show the results when repeating this process over a multiplicity of scenarii. `Dune` and the other merging methods rely on one or multiple clustering results – in this work, clusterings from `SC3`, `Seurat`, and `Monocle`. Because each of these methods have tuning parameters than can affect their performance, we ran each of the three clustering methods on a grid of tuning parameter values for all 4 datasets, as described in the “Methods, Data analysis” section. The AUARIC for the three merging methods across these 36 scenarios are displayed in Figure 3c and Table S2. Overall, `Dune` clearly outperformed the other two merging methods. Table S2 recapitulates all rankings. In particular, in 29 out of the 36 evaluations, `Dune` resulted in the highest ARI increase and was the lowest performer only twice.

### Dune outperforms other methods in terms of the resolution-replicability trade-off

We then considered the replicability of the clusters found by `Dune` compared to the other two merging strategies. We measured replicability by evaluating whether the method finds similar clusters for multiple independent datasets – for example, datasets on the same biological system but from different labs or technologies. We considered two pairs of datasets: The two mouse brain AIBS Smart datasets from the Allen Institute and the two human pancreas datasets **Baron** and **Segerstople**. To measure replicability, we relied on the `MetaNeighbor` algorithm from Crow et al. [22], which identifies replicable clusters between pairs of datasets (see “Methods, metaneighbour” for description). The replicability of a set of clusters was then defined as the fraction of cells in replicable clusters. We used this measure to compare `Dune` to other merging procedures.

#### Illustration of the trade-off between resolution and replicability

Figure 4a displays replicability vs. resolution for a wide range of clustering results, where three clustering methods (`SC3`, `Seurat`, and `Monocle`) were run with a large grid of tuning parameter values, on the pair of mouse brain datasets. This clearly demonstrates the trade-off between replicability and resolution: As the number of clusters increased, the fraction of cells in replicable clusters decreased, regardless of the clustering method used. While the actual trade-off is specific to the biological context and the pair of datasets that are being considered, it should be noted that a similar trade-off is clearly visible when applying the same type of analysis to the human pancreas datasets (Figure S2). Note that although it might be tempting to use this figure to contrast and benchmark clustering methods, this would not appropriate. Indeed, pre-processing steps were not identical between the three methods – as described in “Methods, Data analysis” – and, as such, no direct comparison is possible.

**Figure 4:**
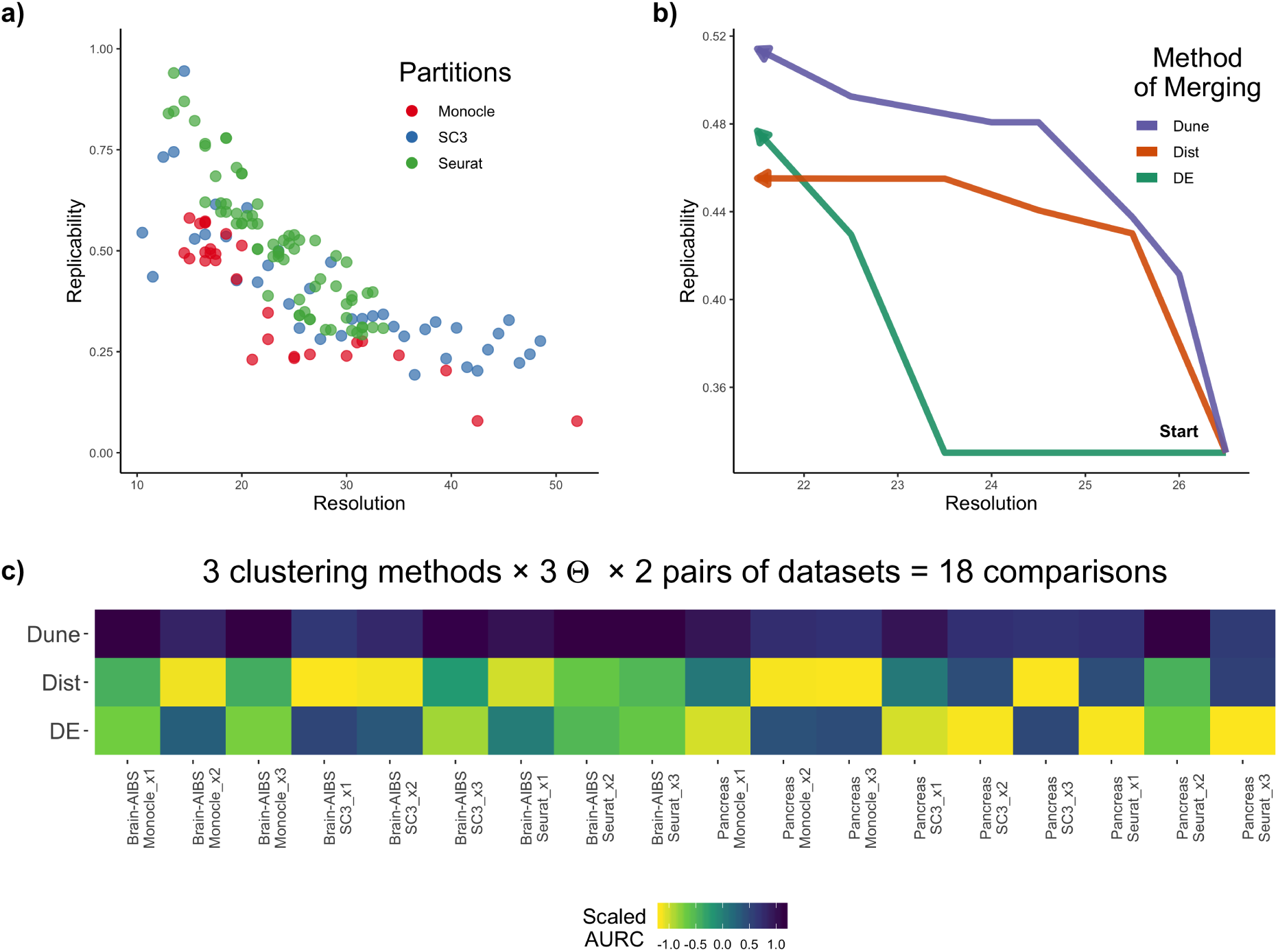
Dune correctly navigates the resolution-replicability trade-off. Panel **a.** `SC3`, `Seurat`, and `Monocle` were run on the two AIBS snRNA-Smart datasets, as described in Methods, for a wide range of tuning parameter values. Then, the `MetaNeighbor` method was used to find the clusters that replicate between these two datasets. Replicability was then computed as the fraction of cells in replicable clusters. There is an apparent trade-off between resolution and replicability. Panel **b.** For a given point from **a**, we merged the clusters and tracked how replicability evolved as we decreased resolution. Panel **c.** For each of the curves in **b**, we computed an area under the replicability curve (AURC). This was repeated over the three clustering methods, each with three different values of their respective tuning parameter *θ*, and for the two pairs of datasets. AURC were scaled column-wise for display in the pseudocolor image.

#### Comparison of merging methods

As pairs of clusters are merged, the resolution decreases, so a well-performing merging method is one that improves the replicability of the clusters. Therefore, a natural way to benchmark merging methods is to measure how and if replicability improves as the number of clusters is reduced. For example, in Figure 4b, `Seurat` was run with *θ*_*Seurat*_ = 1.7 on each of the two AIBS Smart datasets. The two sets of clusters were then merged using the three different merging methods, independently on each dataset. `Dune` also used the clusterings from `SC3` (*θ*_*SC*3_ = 15) and `Monocle` (*θ*_*Monocle*_ = 15). At each step of the merging, we then tracked how replicability evolves. All three merging methods outputted sets of clusters with increasing replicability as resolution decreases, but `Dune` produced clusters that have higher replicability compared to the other two. The area under the replicability curve (AURC) was computed for each merging method. This was repeated for the three clustering methods, each with three values of their respective tuning parameter *θ*, and two pairs of datasets, which lead to 18 comparisons, depicted in the pseudocolor image of Figure 4c. `Dune` outperformed the other two merging methods in all 18 comparisons. Note that, as in the previous section, merging for the other methods was stopped at the resolution level where `Dune` stopped, which provided these methods with more information than they would otherwise have had.

### Dune has a natural stopping point

Unlike other merging methods, `Dune` provides a meaningful stopping point, i.e., it keeps merging clusters until no improvement in average ARI occurs. By contrast, the two hierarchical merging methods continue to merge until there is only one cluster, which is not biologically meaningful or interesting.

Each clustering method has some strengths and drawbacks: `Dune`’s stopping point identifies the level of resolution where all clustering algorithms are close to full agreement. Furthermore, at the stopping point, the clusters overlap very well with gold-standard clusters. In Figure S3a, the outputs from `SC3`, `Seurat`, and `Monocle` were used as inputs to `Dune` on the AIBS snRNA-Smart dataset. After merging with `Dune`, the clusters from `SC3` overlap well with the Allen Institute subclass labels. Indeed, the ARI between the `SC3` clusters and the subclasses increases from ∼ .63 before merging to ∼ .83 after merging.

### Dune robustness analysis

#### Robustness to poor clustering inputs

Since `Dune` takes as input the results from clustering algorithms, it is sensitive to the quality of the clusterings produced by these algorithms. In general, `Dune` will not be able to produce good clusters when merging only clusters that capture no underlying biological signal. However, we showed that `Dune` is robust to a mix of “good” clustering inputs and “bad” clustering inputs. We used as “good” inputs the results of `SC3`, `Seurat`, and `Monocle` and as “bad” inputs fully random clusters (see the “Methods, Data analysis” section). Then, the replicability of the “good” clusterings was measured as merging happened and the AURC was computed and compared to the AURC when there was no “bad” inputs. As more and more “bad” clusters were added (Figure S3b), `Dune` still improved the replicability of the “good” clusters as it merged them, even when half of the clusters used as inputs were random. Hence, `Dune` can recover from very poor clustering inputs.

#### Robustness to sample size

We investigated how `Dune` handles datasets with an ever-smaller number of cells. To simulate such datasets, we downsampled the two pancreas datasets. Downsampling could affect both the quality of input clusters and the merging procedure of `Dune`. To disentangle these two effects, we downsampled the two human pancreas datasets after running `SC3`, `Seurat`, and `Monocle`, but before running `Dune`. We then measured how and whether merging still improved the cluster replicability by computing the AURC and constrasting it to its value without downsampling (see the “Methods, Data analysis” section for more details).

When the datasets were downsampled to between 90% and as low as 10% of the original number of cells, `Dune` still correctly navigated the trade-off between resolution and replicability (Fig. S3c). Only when fewer than 10% of the cells were used (which amounts to datasets of fewer than 200 cells) did `Dune`’s capacity to improve cluster replicability worsen noticeably. This demonstrates that the method is very stable to the number of cells.

## Discussion

We have introduced `Dune`, a new method for navigating the resolution-replicability trade-off in cluster analysis and for aggregating clustering results from multiple algorithms. We stress that `Dune` is not a new clustering algorithm; instead, it relies on different clustering methods to identify the highest resolution at which cluster quality (i.e., replicability across datasets) remains high. In doing so, `Dune` identifies the commonalities of the input clusterings and uses this to improve each of these clusterings. The method is stable with respect to the quality of the input clusterings as well as to the number of cells/observations to be clustered. Furthermore, as a result of merging clusters, `Dune` provides a sensible hierarchy on the clusters based on their commonality across different methods. As we go up in this hierarchy, the number of clusters is reduced, but their replicability improves. In this regard, `Dune` outperforms more commonly used hierarchical merging methods.

`Dune` automatically stops at a meaningful resolution level, where all clustering algorithms are in agreement, while the other methods either keep merging until all clusters are merged into one or require user supervision to stop early. This feature helps users in identifying reliable structure in their scRNA and snRNA datasets. The manual choice of a stopping point is difficult since, in practice, it is often impossible to measure replicability given the lack of a second appropriate dataset.

`Dune` relies on the adjusted Rand index (ARI) to decide which clusters to merge. Because of this, it currently cannot be used with clustering methods that do not cluster all cells unambiguously, e.g., with soft or fuzzy clustering methods which could assign some cells to multiple clusters based on weights. Other approaches, such as `RSEC`, leave some cells unclustered. For now, using such methods as input to `Dune` would require forcing a hard assignments of the cells (possibly to their nearest cluster) or excluding ambiguous/unclustered cells. Extensions of the ARI to fuzzy clustering have been proposed [23, 24] and would need to be evaluated.

This manuscript focuses on the question of unsupervised clustering. Recent work in supervised clustering [25–28] has proposed labeling cells in a new dataset by relying on information contained in other datasets or even cell atlases. In practice, these methods define marker genes for known cell types and build classifiers to assign new cells to these cell types. In particular, `Garnett` [29] allows a hierarchical clustering structure, but one that needs to be predefined, and `scClassify` [30] uses the `HOPACH` [31] algorithm to establish a hierarchy in the training dataset. Most of these algorithms can also identify new cell types not present in the reference. It is therefore possible to use a supervised clustering method to identify the cells of a dataset that have a known cell types. If these cells do not provide information to help cluster the rest of the cells, we can remove them, and then use unsupervised clustering methods and `Dune` on the remaining cells.

While the method we propose has only been benchmarked on scRNA-Seq and snRNA-Seq datasets, it is a general framework that can be applied to any clustering setting.

**Table 1:** Adjusted Rand index. Contingency table for two partitions **P**_1_ and **P**_2_.

| | $\mathcal{C}_1^2$ | $\mathcal{C}_2^2$ | … | $\mathcal{C}_{k_2}^2$ | Sums |
| --- | --- | --- | --- | --- | --- |
| $\mathcal{C}_1^1$ | $n_{1,1}$ | $n_{1,2}$ | … | $n_{1,k_2}$ | $a_1$ |
| $\mathcal{C}_2^1$ | $n_{2,1}$ | $n_{2,2}$ | … | $n_{2,k_2}$ | $a_2$ |
| ⋮ | ⋮ | ⋮ | $\ddots$ | ⋮ | ⋮ |
| $\mathcal{C}_{k_1}^1$ | $n_{k_1,1}$ | $n_{k_1,2}$ | … | $n_{k_1,k_2}$ | $a_{k_1}$ |
| Sums | $b_1$ | $b_2$ | … | $b_{k_2}$ | |

## Acknowledgments

We thank Quentin Dumont and Frank Herbert [32] for inspiration for the `Dune` name.

This work used the Extreme Science and Engineering Discovery Environment or XSEDE (which is supported by National Science Foundation grant number ACI-1548562) PSC, Bridges Regular and Large Memory at the Pittsburgh Supercomputing Center through allocations TG-IBN180019 and TG-IBN190010. This work was supported by NIH grant U19MH114830 (JN), U19MH114821 (JG) and R01MH113005(JG). KVdB is a postdoctoral fellow of the Belgian American Educational Foundation (BAEF) and is supported by the Research Foundation Flanders (FWO), grant 1246220N.

## Authors’ contributions

HRB, KS, JN, EP, and SD conceived and designed the study. HRB and KS developed and implemented the method. HRB, KS, SF, and DR analyzed the data. RC provided resources. HRB and SF wrote the initial draft of the manuscript, and KS, KVdB, RC, DR, JG, JN, EP, and SD contributed to revisions.

## Data availability

The Pancreas datasets were downloaded from the Hemberg group website: https://hemberg-lab.github.io/scRNA.seq.datasets/human/pancreas/ on October 1^*st*^, 2018. The AIBS datasets can be obtained from Neuroscience Multi-omics Archive (*RRID* : *SCR_*002001; nemoarchive.org), (*Zeng sn SSv4* https://assets.nemoarchive.org/dat-k7p82j4 and *Zeng sc SSv4* https://assets.nemoarchive.org/dat-55mowp9).

## Code availability

The results from this paper can be reproduced using code from the following GitHub repository: https://github.com/HectorRDB/Dune_Paper. The `Dune` method is implemented in an open-source `R` package available on Github: https://github.com/HectorRDB/Dune *(RRID* : *SCR_*018218) and to be released through the Bioconductor Project (http://www.bioconductor.org).

## Methods

Consider a – possibly high-dimensional – dataset of *n* observations, **X** = {*x*_1_, …, *x*_*n*_}, where *x*_*i*_ ∈ ℝ^*J*^, *i* = 1, …, *n*. For instance, in scRNA-Seq, *x*_*i*_ corresponds to the *J* gene expression measures (i.e., normalized read counts) of cell *i*. Represent the results of any (non-fuzzy) clustering method as a partition, **P**, which splits the set of *n* observations into *k* disjoint subsets or clusters, {𝒞_1_, …, 𝒞_*k*_}, where: **1)** 𝒞_*i*_ ∩ 𝒞_*j*_ = ø, ∀*i, j* ∈ {1, …, *k*}, and **2)** ∪_*i*∈{1,…,*k*}_𝒞_*i*_ = **X**. Accordingly, a collection of *R* clustering results may be represented as multiple partitions, **P**_1_, …, **P**_*R*_, with partition **P**_*r*_ containing *k*_*r*_ clusters, *r* = 1, …, *R*. For each observation *x*_*i*_, denote by 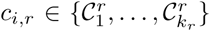 the cluster to which it belongs in partition **P**_*r*_.

The focus of the present manuscript is to develop a general approach to combine clusters within the different partitions, **P**_1_, …, **P**_*R*_, in order to balance the trade-off between cluster resolution and replicability. In the remainder of this section, we first present the Rand index, a well-known measure of concordance between two partitions, and its adjusted version. We also review popular clustering methods in the scRNA-Seq literature and alternative approaches to merge clusters. Finally, we formalize the two key notions of cluster resolution and cluster replicability.

### Adjusted Rand index

The Rand index [16] measures the concordance between two partitions **P**_1_ and **P**_2_. Denote by *a* = |{(*x*_*i*_, *x*_*j*_) ∈ **X**^2^|(*c*_*i*,1_ = *c*_*j*,1_)&(*c*_*i*,2_ = *c*_*j*,2_)}| the number of pairs of observations that are in the same cluster for both partitions **P**_1_ and **P**_2_ and by *b* = |{(*x*_*i*_, *x*_*j*_) ∈ **X**^2^|(*c*_*i*,1_ ≠ *c*_*j*,1_)&(*c*_*i*,2_ ≠ *c*_*j*,2_)}| the number of pairs of observations that are in different clusters for both partitions **P**_1_ and **P**_2_. The Rand index is then the ratio of *a* + *b* over the total number of pairs of observations

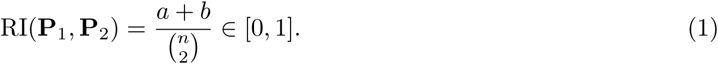

Thus, intuitively, the Rand index is the proportion of pairs of observations for which the two partitions are in agreement.

However, the Rand index does not account for the fact that a pair of observations might be in the same (different) cluster(s) in the two partitions purely by chance. The adjusted Rand index (ARI) [17] adjusts for the level of concordance expected by chance, yielding a value between −1 and +1. Specifically, considering **P** a fixed partition and **R** a random permutation of **P**, then 𝔼[ARI(**P, R**)] = 0, where the expected value is over all cluster permutations (i.e., permutations of the cluster assignments of the observations, while keeping the number of clusters and the sizes of the clusters fixed). Negative values indicate less than the expected level of concordance and positive values indicate more than the expected level of concordance. The ARI relies on the contingency table of two partitions **P**_1_ and **P**_2_, with the (*i, j*)^th^ entry *n*_*i,j*_ defined as the number of observations both in cluster *i* of partition **P**_1_ and cluster *j* of partition **P**_2_ (Table 1). Examples of contingency tables between two partitions can be found in Figures 1a, 1b, 2a, and 2d.

Given the contingency table notation, the adjusted Rand index is defined as

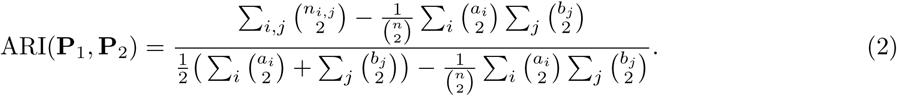

For *R* partitions, the level of concordance can be quantified by the average ARI for all possible pairs of partitions

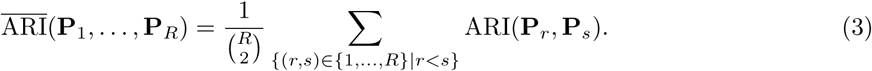

Note that, in the case of *R* = 2 partitions, this is simply the ARI between the two partitions. If one considers the matrix of pairwise ARIs between partitions, such as displayed in Figures 2b and e, then the average ARI is defined as the mean of the upper(or lower)-triangular matrix.

### ARI merging with Dune

Given *R* partitions (possibly the result of different clustering algorithms or different tuning parameter values for the same clustering algorithm or both), **P**_1_, …, **P**_*R*_, with **P**_*r*_ containing *k*_*r*_ clusters, *r* = 1, …, *R*, `Dune` seeks to improve the overall agreement among these, as measured by the average ARI, through an iterative process of merging clusters within partitions.

Specifically, `Dune` searches over each partition **P**_*r*_ and over each of 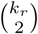 pairs of clusters in **P**_*r*_ for the pair which produces the largest improvement in ARI when merged, i.e.,

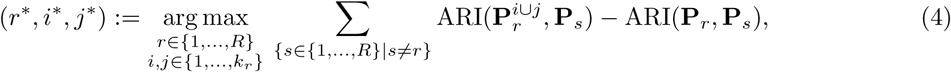

where 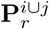 is the partition created by merging clusters 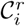 and 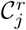 in partition **P**_*r*_

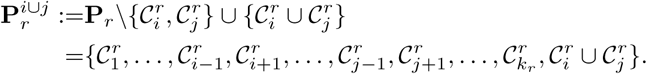

`Dune` amounts to a greedy algorithm for maximizing the average ARI, 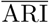. At each step, we find the pair of clusters that, when merged, lead to the greatest improvement in 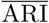. Once we have identified this pair of clusters, we update the collection of partitions: 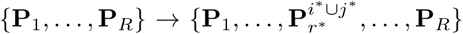. We continue iterating until no beneficial merge can be identified, that is, we stop updating when

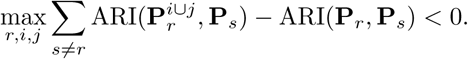

This greedy approach means that each update step is constrained to merging a single pair of clusters from a single partition. As such, we never merge three clusters together in one iteration or two pairs of clusters in the same or in separate partitions. This ensures that, in our applications, we do not converge to the naive optimal solution of merging all clusters, which does represent a full agreement between the partitions but is of no practical interest.

While `Dune` provides a natural stopping point for merging, it is also possible to stop earlier in the merging process, by tuning the merging parameter *m*_Dune_, which is defined as the fraction of ARI improvement over the total ARI improvement. For example, *m*_Dune_ = .5 means that `Dune` returns the merged partitions that have a mean ARI halfway between the mean ARI of the original partitions and the mean ARI of the final ones.

### Computational implementation and run time

The `Dune` algorithm has been implemented in an open-source `R` package available on Github: https://github.com/HectorRDB/Dune. It is implemented in a fully-parallel and efficient manner. Run time for a large dataset of ∼ 100, 000 cells with 3 partitions is under 15 minutes with 10 CPUs. The package also contains plotting functions used to create many panels of the paper, as well as options to create GIFs and track the evolution of mean ARI or confusion matrices across the merging steps.

### Clustering algorithms for scRNA-Seq data

Any combination of clustering algorithms and associated tuning parameters, applied to an appropriate dataset, can produce a set of partitions that can be used as input to `Dune`. However, as our work was motivated by the classification of cells based on transcriptomic signatures, we will focus on this particular setting to benchmark `Dune`.

In the descriptions below, we use the notation from the original papers to describe the tuning parameters of each method; the same notation may therefore correspond to different parameters depending on the algorithm.

`SC3` [2] is a consensus clustering method that involves performing *k*-means clustering on different dimensionality reductions of the input dataset. A hierarchical clustering method is then applied to the resulting consensus matrix. The main parameter is the number of clusters *k*, which is used both in *k*-means and to cut the hierarchical clustering tree. The method provides an estimate of the optimal value of this parameter, *k*_0_, based on the number of eigenvalues of the centered and scaled distance matrix that are significantly different from 0 (see Kiselev et al. [2] for more details). For large datasets, there exists a hybrid version of the algorithm, where the full `SC3` clustering method is run on only a fraction of the cells to identify the clusters and the rest of the cells are assigned to the clusters using a support vector machine (SVM) algorithm.

`Seurat`’s clustering algorithm (*SEURAT, RRID* : *SCR_*007322) has evolved over the different versions of the software; here, we focus on version 3 [3] (we specifically use version 3.1.1). The algorithm first reduces the dimension of the data by selecting the first *p* principal components (PCs) and then computes a *k*-nearest neighbor (*k*-NN) graph. After refining the graph, it groups cells together using, as default, the Louvain algorithm [33]. The two main tuning parameters are the number of neighbors *k* used to build the *k*-NN graph and the resolution parameter for the Louvain algorithm.

`Monocle`’s clustering algorithm has also changed and we focus on version 3 [4] (implemented in the `Monocle3` package, although we keep the name `Monocle` for simplicity; we specifically use version 0.1.3). `Monocle`’s clustering algorithm is similar to the one implemented in `Seurat`, with a few differences. After initial dimensionality reduction based on principal component analysis (PCA), `Monocle` performs another dimensionality reduction step using uniform manifold approximation and projection (UMAP) [34, 35] and relies on that representation to build the *k*-NN graph. It then clusters cells using, by default, the Leiden algorithm [36].

Resampling-based sequential ensemble clustering (`RSEC` [7]) is a consensus method over user-supplied clustering algorithms and their associated tuning parameters. In order to improve the stability and tightness of the clusters, it also provides the option to perform clustering on subsamples of the observations, as well as sequential clustering. However, in this paper, we mainly use `RSEC` for its final step of hierarchical merging, see section Existing methods to merge clusters.

#### Method parameters

For each method, we only tune the main parameter. For `Seurat`, however, there are two main tuning parameters. The *k* parameter controls the number of neighbors used to build the *k*-NN graph, while the resolution parameter defines the neighborhood in the Louvain clustering algorithm. In practice, the *k* parameter has much less impact than the resolution parameter (see Figure S1). Moreover, depending on the value of the resolution, increasing *k* either increases or decreases the final number of clusters. As a result, we only consider changing the resolution parameter.

For ease and generality of notation, we will denote each method’s main tuning parameter by *θ* and define *θ* such that increasing *θ* increases the number of clusters. Thus, for the methods described above, *θ*_*SC*3_ = *k, θ*_*Seurat*_ = Resolution, and *θ*_*Monocle*_ = −*k*. Each combination Θ = {*θ*_*SC*3_, *θ*_*Seurat*_, *θ*_*Monocle*_} of the three parameters defines a set of partitions that serves as input for `Dune`.

### Existing methods to merge clusters

Once a set of clusters has been identified, one can build a hierarchical tree for these clusters and then merge clusters that are similar. This involves specifying a measure of distance or similarity between individual observations (i.e., cells) as well as between clusters. It should be noted that the distance used to build the tree of clusters need not be the same as the distance used to merge clusters.

For scRNA-Seq datasets, commonly used between-cell distance measures include the Euclidean distance and one minus the Spearman correlation coefficient. Between-cluster distances include classical linkage measures used in hierarchical clustering, e.g., maximum/minimum/average of all pairwise distances between observations in two clusters or distance between the cluster averages or medoids. For scRNA-Seq, another sensible between-cluster distance measure is the proportion of differentially expressed (DE) genes between clusters [7, 8]. A detailed discussion of such measures is out of the scope of this manuscript[37].

Here, we consider two possible ways of merging. In both cases, we compute the cluster medoids (median of the cluster) based on the log-transformed count matrix (adding 1 to avoid taking the log of zero). We then build a hierarchical tree of clusters using the Euclidean distance between the cluster medoids. The first merging approach directly uses this tree to decide how to merge clusters. Specifically, clusters are merged bottom-up, starting with the two clusters that are closest in the tree and then iteratively until all clusters are merged. The parameter *m*_*Dist*_ = *n*_*merges*_, the number of merges (between 0 and the initial number of clusters minus one), controls the amount of merging. The second approach follows the method implemented in `RSEC`. It computes the percentage of DE genes between clusters, using the `limma` package [20] (*LIMMA, RRID* : *SCR_*010943), where a gene is declared DE if its nominal FDR adjusted *p*-value is below 0.05 [21]. The main tunable parameter is *m*_*DE*_ = *α* ∈ [0, 1], the threshold for the percentage of DE genes below which we merge. We name these two methods `Dist` and `DE`, respectively.

### Cluster replicability using MetaNeighbor

We quantify the replicability of clusters across datasets by applying a modified version of unsupervised MetaNeighbor [22] (*MetaNeighbor, RRID* : *SCR_*016727). MetaNeighbor requires as input a set of unnormalized datasets, a set of cluster labels, and a set of highly variable genes. It uses a cross-dataset validation scheme to quantify how well clusters match across datasets. Given any two datasets, MetaNeighbor builds a cell-cell similarity network based on the Spearman correlation over the set of highly variable genes. One of the datasets is treated as a test dataset, where all cluster labels are hidden, the other dataset is treated as a training dataset, whose labels are propagated to the test dataset through the cell-cell similarity network. Each pair of clusters (one in the training dataset, the other in the test dataset) receives a score based on how well the training cluster predicts the labels from the test cluster. This score is the area under the receiver operator characteristic curve (one-vs-one AUROC). We define the best matching cluster as the test cluster which dominates all other test clusters (one-vs-one AUROC > 0.5). Finally, we reduce the test set to the two best matching clusters, recompute an AUROC, which we call one-vs-best AUROC, and record this as the pair’s final score. Then the role of the test and training datasets are reversed. A cluster is considered replicable if there is a cluster in the other dataset such that the clusters are reciprocal best hits with a high AUROC score (one-vs-best AUROC > 0.6 both ways). See Crow et al. [22] for details.

The **replicability score of a cluster** is defined as the fraction of cells contained in replicable clusters. More specifically, for a comparison of two datasets, we enumerate replicable clusters in each dataset, then deduce the number of cells that are in replicable clusters, sum this number across datasets, and divide by the total number of cells.

We used MetaNeighbor’s `variableGenes` procedure to select genes that were detected as highly variable across all datasets. For performance reasons, the `variableGenes` procedure was applied to a random subset of 50,000 cells for datasets exceeding that size. However, the full datasets were use for the rest of the analysis. In the end, we obtained a set of 541 highly variable genes for the Allen brain datasets and 2, 147 genes for the pancreas datasets.

### Case studies

#### AIBS Smart mouse brain datasets

We used the two AIBS Smart datasets produced as part of the Brain Initiative Cell Census Network (BICCN: *RRID* : *SCR_*015820) and described in Yao et al. [11], one is single-cell (*Zeng sn SSv4* https://assets.nemoarchive.org/dat-k7p82j4) and the other is single-nuclei (*Zeng sc SSv4* https://assets.nemoarchive.org/dat-55mowp9). We use the subclass labels as gold-standard cluster labels for these datasets. Those datasets can be downloaded from the Neuroscience Multi-omics Archive (*RRID* : *SCR_*002001; nemoarchive.org). More details on the parent data set (https://assets.nemoarchive.org/dat-ch1nqb7) and data access can be found in Yao et al. [11].

#### Human pancreas datasets

We focus on two datasets from [18] (8, 568 cells) and [19] (3, 514 cells) which we name **Baron** and **Segerstople**, respectively. Both datasets were downloaded from the https://hemberg-lab.github.io/scRNA.seq.datasets/ on October 1^*st*^, 2018. We use the clusters from the original publications as gold-standard cluster labels.

### Data analysis

Except when otherwise specified, all methods and algorithms were run with default parameters or, if no available default, with the parameters recommended in the vignette or tutorial.

#### Pre-processing

Count matrices were filtered to remove lowly-expressed genes with fewer than *i* reads in *j* cells. See Table S1 for values of *i* and *j* for each dataset.

As indicated below, we follow different normalization strategies before running `Seurat` and `Monocle` in order to obtain more diverse clustering results. This is appropriate, as the goal of the manuscript is not to compare different clustering methods, but rather different merging methods for given clustering results. The merging methods that `Dune` is compared to rely on only one clustering input; we therefore seek to benchmark merging methods using a variety of clustering inputs.

#### Seurat

Following the tutorial, we run `FindVariableFeatures` and `ScaleData` to normalize the data. Counts are log-transformed (adding 1 to avoid taking the log of zero) and normalized by sequencing depth. For the two pancreas datasets, batches are also normalized using the `scaleData` function. Following principal component analysis, `FindNeighbors` and `FindClusters` are run for a number of neighbors *k* in {30, 50, 70} and resolution *θ* from 0.3 to 2.5 in increments of 0.1

#### SC3

The algorithm is run on a dataset normalized as above with the `Seurat` pipeline. The optimal value of *k, k*_0_, is computed using the `sc3_estimate_k` function. The parameter *θ* is transformed to be *θ*_*SC*3_ = *k* − *k*_0_. `SC3` is then run for values of *θ* ranging from −15 to +15.

#### Monocle

`zinbwave` [7] is first used for normalization and dimensionality reduction on the filtered count data. For the two pancreas datasets, batches are included as model covariates. We select *K*, the number of reduced dimensions, based on a visual representation for each dataset, see Table S1. This first step of dimensionality reduction is followed by another using UMAP [35] with two dimensions. The resulting two-dimensional representation is then used to build the *k*-NN graph, with *k* ranging from 10 to 150 in increments of 10.

#### Dune

For a given set of values for Θ = {*θ*_*SC*3_, *θ*_*Seurat*_, *θ*_*Monocle*_}, we get three sets of cluster labels that we can use as input to `Dune`.

#### Building the hierarchical tree

The output of each clustering method is used as input to `RSEC`’s `makeDendogram` function. Then, we either cut the tree using `R`’s `cutree` function or `RSEC`’s `mergeClusters` function.

#### Producing “bad” clusters

For each value of the tuning parameters Θ, on the pancreas datasets, we add fully random inputs to `Dune`. That is, we create “bad” clusterings by randomly assigning each cell a number (or cluster label) between 1 and (*k*_*SC*3_ + *k*_*Monocle*_ + *k*_*Seurat*_)*/*3, where *k* denotes the number of clusters for a particular clustering algorithm. Since cells are assigned randomly, the size of the clusters will vary, but all clusters have the same expected size. To account for the stochastic nature of this procedure, we repeat this 10 times.

#### Downsampling

Downsampling the number of cells at the beginning of the analysis pipeline would affect both the quality of the clustering results and the quality of the merging with `Dune`. As such, to test only the stability of `Dune` to the number of cells, we downsample the cells just before running `Dune`, that is, the clustering algorithms are run on the full dataset but only a subset of the dataset is used to decide which clusters to merge and in which order. Afterwards, cells that are not in the subsample are assigned to the merged clusters based on their original cluster labels. That is, if Cluster 1 and 2 are merged, all cells that were originally in Cluster 1 and 2, even those not selected in the downsampling and used as input to `Dune`, are assigned to the merged cluster.

Most of the code was run using xsede [38].

## Supplementary Material

### Supplementary methods

**Table S1:** Parameters for processing the datasets. Each dataset is filtered such that we keep all genes with a least *i* reads in *j* samples. Then, `zinbwave` is run with *K* dimensions.

| Dataset | $i$ | $j$ | $K$ |
| --- | --- | --- | --- |
| AIBS scRNA-Smart | 50 | 50 | 30 |
| AIBS snRNA-Smart | 50 | 50 | 14 |
| Baron | 5 | 5 | 10 |
| Segerstople | 5 | 5 | 20 |

**Figure S1:**
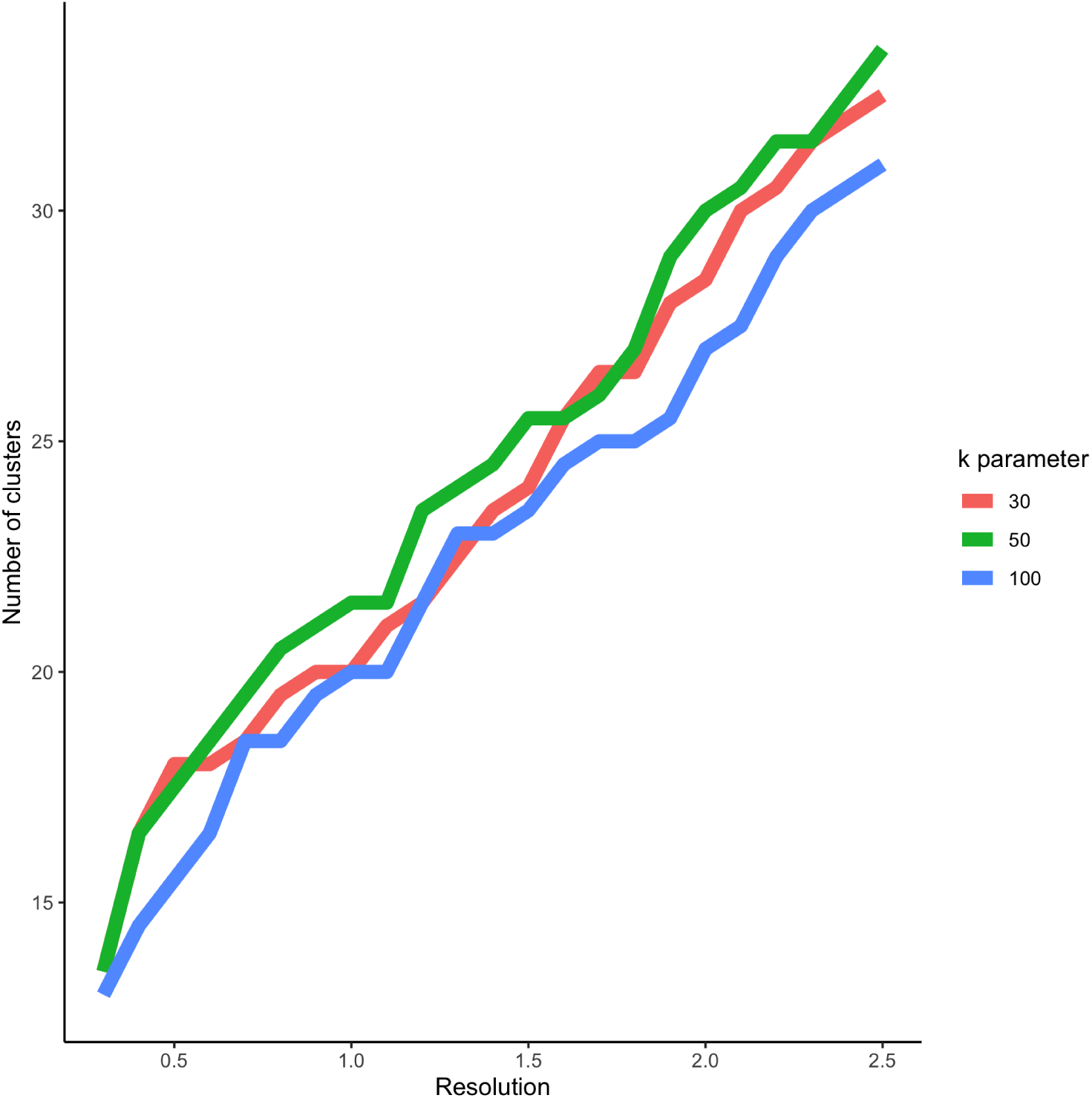
Impact of Seurat’s two main tuning parameters on the number of clusters. The `Seurat` algorithm is run on the two AIBS snRNA-Smart datasets, for a grid of tuning parameter values. The average number of clusters found in both datasets is then computed. For increasing values of the resolution parameter and fixed values of the *k* parameter, the number of clusters is always increasing. On the other hand, for increasing values of the *k* parameter and fixed values of the resolution parameter, the number of clusters can either increase or decrease. This can be seen in the fact that the curves are all increasing but intersect multiple times.

### Supplementary results

**Table S2:** Ranking of merging methods over all 36 comparisons for improving ARI with gold standard. See Figure 3.

|  | 1 | 2 | 3 |
| --- | --- | --- | --- |
| DE | 2 | 21 | 13 |
| Dist | 5 | 10 | 21 |
| Dune | 29 | 5 | 2 |

**Figure S2:**
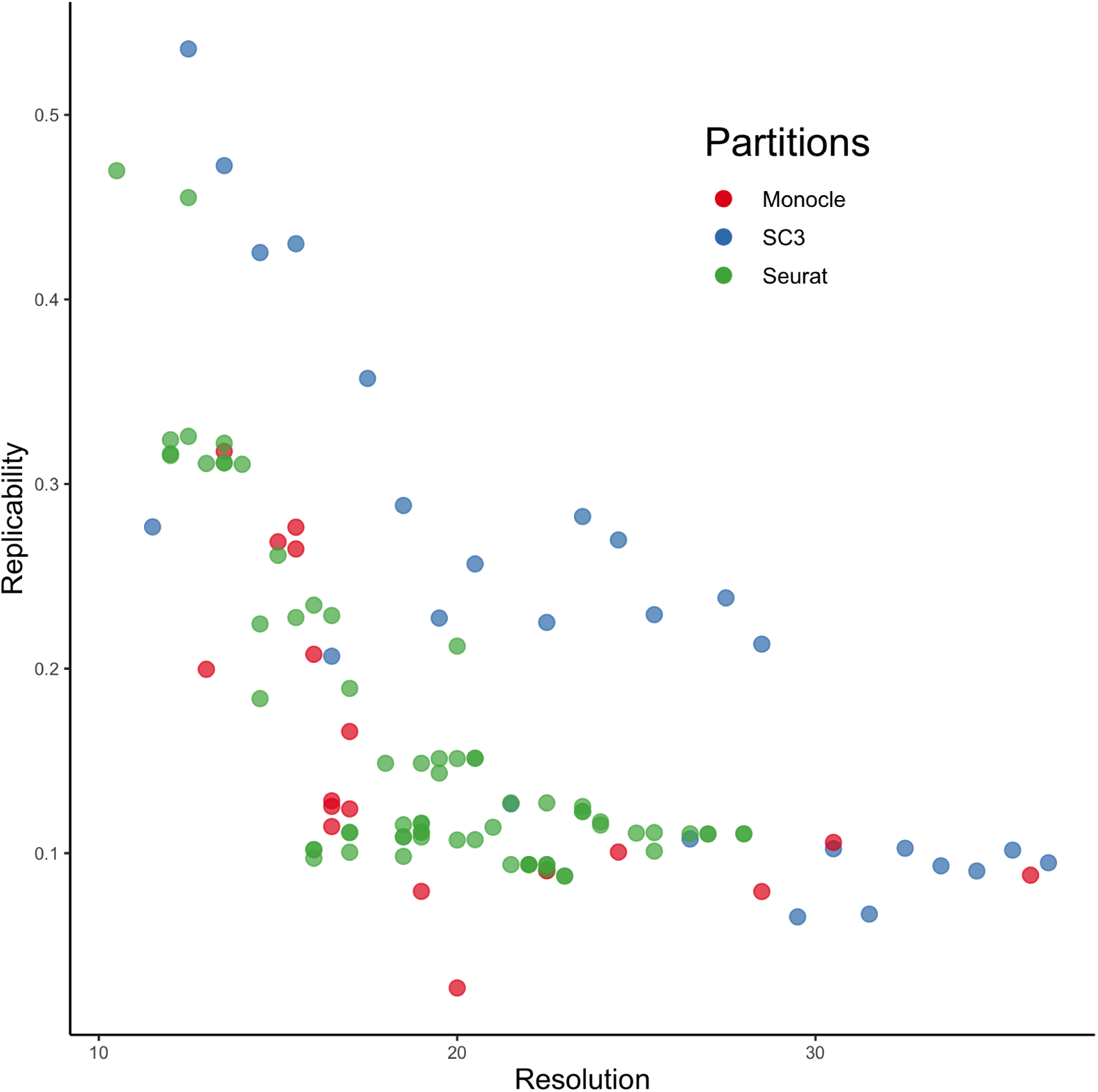
Resolution-replicability trade-off on the Pancreas datasets. `Seurat`, `SC3`, and `Monocle` are run on the two Pancreas datasets, as described in Methods, for a wide range of tuning parameter values. Then, the `MetaNeighbor` method is used to compute replicability scores for the resulting clusters between these two datasets. An apparent trade-off between replicability and resolution is visible.

**Figure S3:**
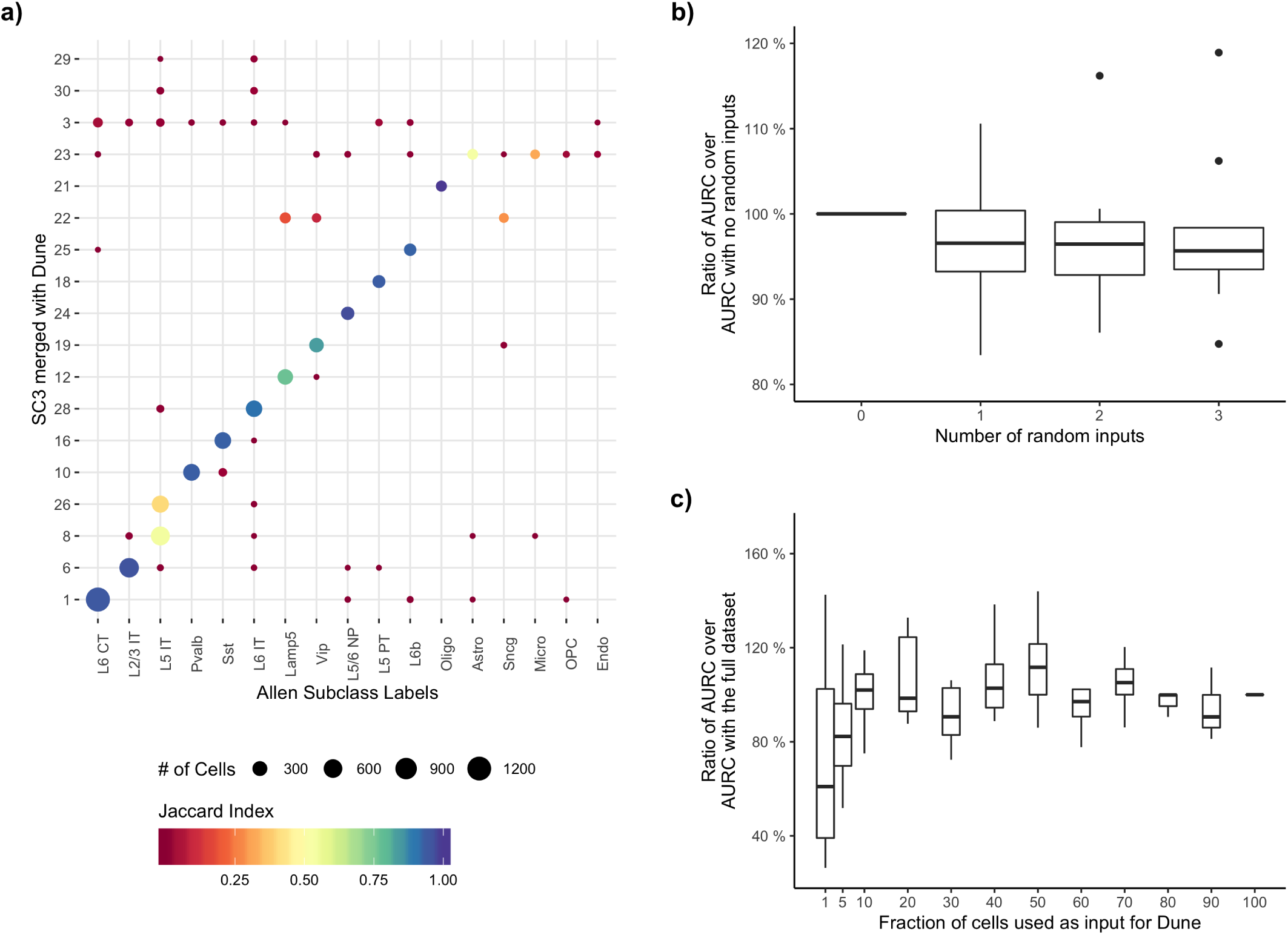
Dune robustness analysis. Panel **a.** Fully merging `SC3` with `Dune` produces meaningful highlevel biological clusters, as can be seen by the overlap between the clustering and the Allen subclass labels. Panel **b.** Adding an increasing number of random clustering inputs to `Dune` impacts only slightly the resolution-replicability area under the curve when merging the other correct clusters. Panel **c.** Likewise, `Dune` is stable to decreasing the number of input cells, as low as 10% of the original sample size.

